# A multidimensional assessment of in-host fitness costs of drug resistance in the opportunistic fungal pathogen *Candida glabrata*

**DOI:** 10.1101/2023.07.03.547612

**Authors:** Amir Arastehfar, Farnaz Daneshnia, Hrant Hovhannisyan, Nathaly Cabrera, Macit Ilkit, Jigar V. Desai, Toni Gabaldón, Erika Shor, David S. Perlin

**Affiliations:** Center for Discovery and Innovation, Hackensack Meridian Health, Nutley, NJ 07110, USA; Division of Infectious Diseases, Massachusetts General Hospital, Boston, MA 02114 USA; Department of Medicine, Harvard Medical School, Boston, MA 02115 USA; Institute of Biodiversity and Ecosystem Dynamics (IBED), University of Amsterdam, Amsterdam1012 WX, The Netherlands; Life Sciences Programme, Supercomputing Center (BSC-CNS), Barcelona, Spain; Institute for Research in Biomedicine (IRB Barcelona), The Barcelona Institute of Science and Technology, Barcelona, Spain; Division of Mycology, Department of Microbiology, Faculty of Medicine, University of Çukurova, Adana, Turkey; Georgetown University Lombardi Comprehensive Cancer Center, Washington DC 20057, USA; Catalan Institution for Research and Advanced Studies, Barcelona Spain; Centro de Investigación Biomédica en Red de Enfermedades Infecciosas (CIBERINFEC), Barcelona, Spain; Department of Medical Sciences, Hackensack Meridian School of Medicine, Nutley, New Jersey, USA

**Keywords:** *Candida glabrata*, fluconazole resistant, echinocandin resistant, multidrug resistant, fitness cost, intracellular replication, gut colonization, systemic infection

## Abstract

The global rise of antimicrobial resistance poses a serious threat to public health. Because drug-resistant (DR) pathogens typically carry mutations in genes involved in critical cellular functions, they may be less fit under drug-free conditions than their susceptible counterparts. As such, the limited use of antimicrobial drugs has been proposed as a practical strategy to diminish the prevalence of DR strains. However, in many cases the fitness of DR pathogens under host conditions is unknown. *Candida* (*Nakaseomyces*) *glabrata* is a prevalent opportunistic fungal pathogen notable for its high rate of fluconazole resistance (FLZR), echinocandin resistance (ECR), and multidrug resistance (MDR) relative to other *Candida* pathogens. Nonetheless, the fitness of *C. glabrata* MDR isolates is poorly characterized, and studies of FLZR isolate fitness have produced contradictory findings. Two important host niches for *C. glabrata* are macrophages, in which it can survive and proliferate, and the gut. Herein, by employing a comprehensive collection of clinical and isogenic *C. glabrata* isolates, we show that FLZR *C. glabrata* isolates are less fit inside macrophages than susceptible isolates and that this fitness cost is reversed by acquiring ECR mutations in *FKS1/2* genes. Interestingly, dual-RNAseq revealed that macrophages infected with DR isolates mount an inflammatory response whereas the intracellular DR cells downregulate processes required for in-host adaptation. Consistently, DR isolates were outcompeted by their susceptible counterparts in the context of gut colonization and in the kidneys of systemically infected mice, whereas they showed comparable fitness in the spleen. Collectively, our study shows that macrophage-rich organs, such as the spleen, favor the retention of DR isolates, potentially reducing the utility of limited antifungal use to decrease the burden of DR *C. glabrata* in the context of candidemia.

**Author summary:** The rise of multidrug resistant (MDR) strains of fungal pathogens, notably *Candida glabrata*, poses a significant clinical challenge because of the limited number of antifungal drugs available for use. Thus, it is vital to minimize the prevalence of drug resistance in the clinic. Because in some bacterial and fungal species drug resistance is accompanied by a fitness cost, implementation of limited antibiotic or antifungal drug use in the clinic has been suggested as a practical way to favor the spread of susceptible isolates. However, it is not clear whether this strategy can work for MDR *C. glabrata*, as its fitness costs have not been systematically examined, particularly in the context of the host. Herein, we show that MDR *C. glabrata* isolates can replicate within macrophages as well as susceptible isolates, and this result was consistent with gene expression changes in the infected macrophages. In animal models, MDR strains were unfit in the context of the gastrointestinal tract and kidney, but their fitness in the spleen was comparable to that of susceptible strains. Accordingly, the potential of limited antifungal use to reduce the prevalence of MDR strains of *C. glabrata* strongly depends on the host reservoir of infection.

## Introduction

Antimicrobial resistance (AMR) is a leading cause of death worldwide and is regarded as one of the most pressing current medical challenges (1). Because antimicrobial drugs typically target cellular processes critical for growth and virulence, mutations that cause drug resistance may attenuate critical enzymes causing diminished growth rates, lower virulence, or reduced transmission. Therefore, in the absence of drug pressure, these mutants may have lower fitness than drug-sensitive strains, resulting in gradual eradication of resistant mutants and dominance of susceptible counterparts. Accordingly, restriction in the use of antimicrobial drugs has been proposed as a practical strategy to decrease AMR rate in clinical settings (2, 3). Thus far, this strategy has led to contradictory findings, with some clinical centers reporting a significant reduction in the rate of AMR (4–7) but others reporting otherwise (8–10). Thus, determination of the true, in-host fitness cost of drug-resistant mutations may provide important insight into this issue. Historically, *in-vitro* growth rates of individual strains have been used as proxy for fitness and shown to correlate moderately with *in-vivo* fitness assessments. However, competition assays between resistant and susceptible isolates, especially those conducted in a physiologically relevant *in-vivo* setting, provide deeper insights and are a better proxy for their relative fitness of such strains in the host environment (2, 3).

*Candida* (*Nakaseomyces*) *glabrata* is a major human fungal pathogen and the second leading cause of candidemia in many countries (11). *C. glabrata* rapidly develops resistance during antifungal treatment, with numerous studies reporting alarming increases in the prevalence of fluconazole-, echinocandin-, and multidrug-resistance (FLZR, ECR, and MDR, respectively) (12–17). Fluconazole exerts fungistatic activity in *Candida* and targets Erg11, one of the critical proteins involved in ergosterol biosynthesis, whereas echinocandins (caspofungin, micafungin, and anidulafungin) are fungicidal and act by inhibiting the catalytic subunits of β-1,3-glucan synthase, Fks1 and Fks2. Mechanisms underpinning FLZR in *C. glabrata* mainly involve gain-of-function (GOF) mutations in the transcription factor Pdr1, which results in overexpression of efflux pumps, whereas ECR is mainly associated with mutations in two hotspot regions (HS1 and HS2) of Fks1 and Fks2 (18).

Interestingly, studies have shown that FLZR *C. glabrata* isolates are more virulent in the context of systemic infections (19), less effectively phagocytosed by macrophages, and more strongly adherent to epithelial cells compared to susceptible isolates (19, 20). Additionally, susceptible wild-type and laboratory-generated FLZR *Cg* isolates induced comparable virulence when tested in *Galleria* larvae (21). On the other hand, FLZR *C. glabrata* isolates harboring gain of function (GOF) Pdr1 mutations were found to be more susceptible to oxidative stress associated with innate immune cells, and that incubation in H_2_O_2_ resulted in acquisition of secondary suppressor mutations that inactivated Pdr1 activity and restored oxidative stress resistance (22). Accordingly, laboratory generated FLZR *C. glabrata* isolates were found to be both susceptible to H_2_O_2_ and more effectively killed by neutrophils (23). Similarly, FLZR *C. lusitaniae* isolates harboring GOF mutations in Mrr1 are more susceptible to H_2_O_2_ and therefore secondary suppressor mutations arise during the course of infection to enhance fitness by reversing the FLZR phenotype (24). Given such contradictory observations and the high rates of FLZR and MDR *C. glabrata* isolates reported in epidemiological studies, it remains unclear if these drug-resistant phenotypes are associated with an in-host fitness advantage.

To fill this knowledge gap, we used both growth assays and competition experiments to comprehensively examine *in-vitro*, *in-cellulo* (intra-macrophage), and *in-vivo* (mouse models of systemic infection and colonization) fitness costs of both lab-derived and clinical *C. glabrata* isolates displaying various drug susceptibility profiles. We found that FLZR *C. glabrata* isolates were less fit when exposed to H_2_O_2_ and less fit inside macrophages. Interestingly, the intracellular defective fitness of the FLZR isolates was rescued by introducing echinocandin-resistant mutations in the HS regions of the *FKS1* and *FKS2*, and the degree of rescue varied depending on the nature of the *fks* mutation, with MDR isolates carrying *FKS1^R631G^*and *FKS2^S663P^* showing the lowest and highest fitness, respectively. In keeping with our intra-macrophage results, dual-RNAseq results indicated that, at the later stage of infection, the transcriptional responses of macrophages infected with MDR-*FKS2^S663P^*and susceptible isolates clustered together and showed responses different from macrophages infected with FLZR and the MDR-*FKS1^R631G^* isolates. Interestingly, MDR-*FKS2^S663P^ C. glabrata* downregulated major cellular processes associated with in-host adaptation and were outcompeted by susceptible and FLZR isolates both in the gut during colonization and in the kidney following systemic infection. In contrast, MDR-*FKS2^S663P^* strain showed fitness comparable to susceptible and FLZR isolates in the spleen during systemic infection. In total, our study comprehensively assessed the fitness of various drug-resistant variants of *C. glabrata* and showed that, although FLZR and MDR isolates are generally less fit compared to susceptible isolates, their fitness varies depending on the infection niche in which kidney and gut favor the retention of susceptible isolates, whereas spleen is a permissive reservoir for drug-resistant *C. glabrata* variants. Together, these results may help explain the high rate of FLZR and MDR *C. glabrata* isolates in the clinic.

## Results

### Strain collection and characterization

To comprehensively assess the impact of drug resistance on fitness of *C. glabrata,* we evaluated 45 clinical strains, including susceptible (*n*=11), ECR (*n*= 12), FLZR (*n*=12), and MDR (*n*=10) isolates. These isolates were collected from different geographical locations and belonged to various sequence types (ST) (Supplementary Table 1). Additionally, we generated a set of drug-resistant isolates otherwise isogenic to the wild-type parent strain (CBS138) (*n*= 34). Briefly, randomly selected CBS138 colonies (*n*=7) were used to generate FLZR (*n*= 10), ECR (*n*=8), and MDR (*n*= 9) isolates. To generate the FLZR isolates, CBS138 was incubated in RPMI containing fluconazole (32µg/ml) for 48 hours, followed by washing and plating to obtain colonies on drug-free yeast-peptone-dextrose (YPD) agar plates. Colonies showing fluconazole minimum inhibitory concentrations (MIC)≥ 64µg/ml and harboring GOF mutation in *PDR1* were considered as FLZR. Of note, several FLZR strains obtained in this manner lacked *PDR1* mutations, but were petite (i.e., lacked mitochondrial function), which is a well-established cause of FLZR in *C. glabrata* (25, 26). These mutants were followed-up in a separate study. ECR colonies were generated by incubation of CBS138 in RPMI containing 2X MIC of caspofungin (0.125µg/ml), followed by recovery of colonies on YPD agar plates containing micafungin (0.125µg/ml). Colonies showing elevated echinocandin MICs and containing mutations inside or near the HS1 and HS2 regions of *FKS1* and *FKS2* were considered as ECR. MDR isolates were obtained by incubation of FLZR isolates in RPMI containing caspofungin (0.125µg/ml), followed by recovery of echinocandin-resistant colonies, which harbored mutations in the HS1 and HS2 regions of the *FKS1* or *FKS2* (Supplementary Table 2). Antifungal susceptibility testing (AFST) and sequencing of *PDR1* and *FKS1* and *FKS2* HS regions were performed for all clinical and CBS138-derived isolates (Supplementary Tables 1 and 2).

### Variation in fitness patterns of drug-resistant *C. glabrata* strains during *in-vitro* stress

First, we examined the *in vitro* fitness of all our *C. glabrata* strains (both clinical and isogenic isolates, *n*= 79) by measuring their growth under various types of stress, including those that mimic conditions in the host. For instance, we considered that low pH, nutrient deprivation, and oxidative stress (pH5, 0.2% dextrose, and 5mM H_2_O_2_) mimics the macrophage phagosome environment. Other stresses included high temperature (42^°^C), membrane stress (0.02% sodium dodecyl sulfate [SDS]), endoplasmic reticulum stress (ER stress) (tunicamycin 5µg/ml), and osmotic stress (0.5M NaCl). YPD broth containing standard glucose concentration (2%) and set at pH7 was used as the stress-free control. To measure growth rates, OD600 of overnight cultures was adjusted at 0.2 in either stress-free or stress-containing medium and changes in OD600 were monitored kinetically over 15 hours.

Although there was significant individual variation in growth rates between strains, the following trends were observed. In stress-free YPD, no subset of clinical isolates showed a significant fitness cost, but CBS138-derived, isogenic ECR and susceptible isogenic isolates showed a slightly higher growth rate than FLZR and MDR isolates (Figures 1A and 1B), which is similar to previous findings (21). Regardless of the origin (clinical or CBS138-derived), susceptible and ECR isolates showed significantly higher growth rates in “macrophage-like” conditions, whereas FLZR and MDR isolates were more tolerant to ER stress. Additionally, strain origin-specific fitness costs were observed, such as better growth of isogenic ECR and susceptible isolates at 42^°^C and better growth of clinical FLZR and MDR isolates in the presence of SDS (Figures 1A and 1B). Together, this comprehensive *in-vitro* growth analysis revealed that ECR and susceptible isolates show similar fitness patterns and may confer a fitness advantage under macrophage-like conditions, which constitute a major niche for *C. glabrata* during infection (27).

**Figure 1.**
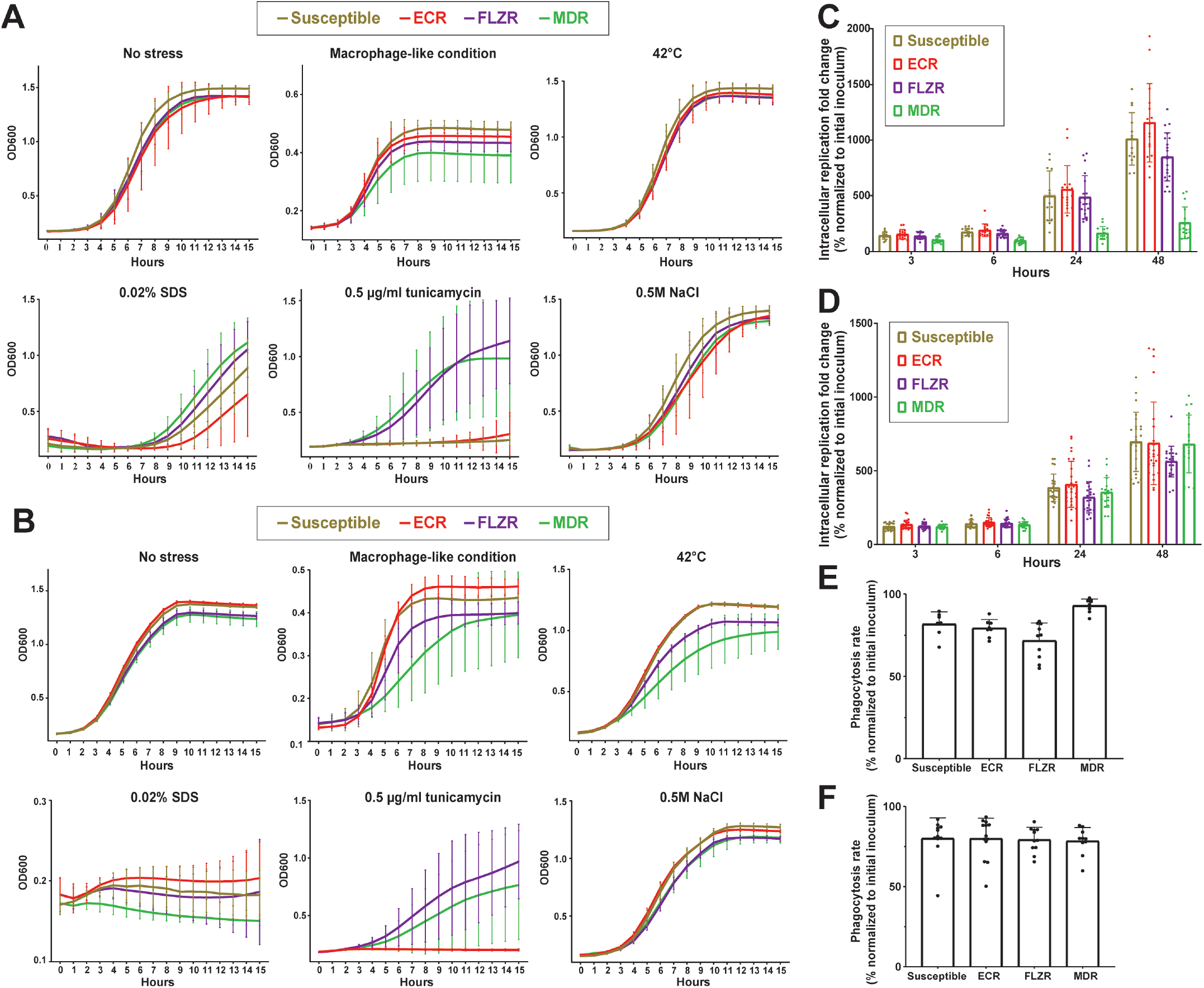
*In vitro* fitness cost assessment of isogenic and clinical *C. glabrata* isolates of various susceptibility profiles. **A.** Clinical susceptible and ECR isolates of *C. glabrata* have relatively high fitness under macrophage-like conditions (see text), whereas FLZR and MDR isolates have relatively high fitness under ER stress. **B.** Isogenic susceptible and ECR isolates of *C. glabrata* have relatively high fitness under macrophage-like conditions, whereas FLZR and MDR isolates have relatively high fitness under ER stress. **C.** In the lab-derived isogenic strain panel, ECR and susceptible isolates had the highest IR rate in macrophages at 48 hrs, whereas FLZR and especially MDR isolates had lower IR rates. **D.** In the clinical isolate strain panel, susceptible, ECR, and MDR isolates had similar IR rates in macrophages, whereas the FLZR isolates had the lowest IR rate. **E.** Isogenic FLZR isolates had the lowest phagocytosis rate. **F.** The four types of clinical isolates were phagocytosed at similar rate. MDR= multidrug resistant, FLZR= fluconazole resistant, ECR= echinocandin resistant.

### ECR and susceptible isolates show high fitness inside macrophages

Given the higher fitness of the ECR and susceptible isolates under “macrophage-like” conditions, we investigated the replication rate of all our *C. glabrata* isolates (*n*= 79) in THP1 macrophages. Macrophages were infected with the multiplicity of infection of 1/1, extensively washed with PBS 3 hrs post-infection (hpi) and treated with fresh RPMI. Intracellular replication (IR) was assessed by plating lysed macrophages at 3-, 6-, 24-, and 48 hpi and counting colony forming units (CFU). Furthermore, phagocytosis rates were determined 3 hpi by plating and CFU counting of the RPMI supernatants.

Interestingly, and consistent with *in-vitro* experiments, CBS138-derived susceptible and ECR isolates showed the highest IR rates, whereas the MDR isolates were the least fit inside the macrophages (Figure 1C). For clinical isolates, the IR rates of the MDR isolates were similar to susceptible and ECR isolates, whereas the FLZR isolates had the lowest IR rates (Figure 1D). This difference between IR rates of clinical and CBS138-derived MDR isolates might reflect the fact that clinical MDR isolates have evolved in the presence of host factors acting as selection pressures, whereas the CBS138-derived MDR strains were derived in an *in-vitro* passaging scheme in the presence of antifungal drugs but absence of host factors, thus lacking any evolutionary pressures for survival in the host.

Consistent with a previous study (20), isogenic CBS138-derived FLZR isolates had the lowest phagocytosis rate (Figure 1E), whereas phagocytosis rates were similar for all clinical isolates regardless of susceptibility profile (Figure 1F). Of note, the lower IR rate of isogenic FLZR isolates was not simply due to their lower phagocytosis rate because, first, clinical FLZR isolates had phagocytosis rates similar to other isolates and, second, all isolates had very similar CFUs at 3 hpi. Altogether, these observations suggested that inside macrophages susceptible, ECR, and clinical MDR isolates have similar fitness, whereas FLZR isolates sustain the highest fitness cost.

### The low intracellular fitness of FLZR isolates can be rescued by certain *fks* mutations

Given that ECR isolates had a significantly higher IR rates than FLZR isolates, we hypothesized that introducing echinocandin resistance into FLZR strains may increase their fitness inside macrophages. However, as described above, we observed that MDR isolates generated by passaging FLZR strains in caspofungin in fact decreased their intra-macrophage fitness. Importantly, these lab-derived MDR isolates had a very narrow range of *fks* mutations, predominated by *FKS2^F659del^*. In contrast, clinical MDR isolates are present other *FKS* hot-spot mutations, such as *FKS1^S629P^*, *FKS2^S663P^,* and *FKS2^F659Y^* (28). Therefore, we used CRISPR-Cas9 to directly introduce specific, clinically relevant mutations in the HS1 of *FKS1* and *FKS2*. Equivalent mutations occurring in Fks1-HS1 and Fks2-HS1, i.e., S629P vs. S663P and R631G vs. R663G (Figure 2A), were introduced into a CBS138-derived FLZR isolate carrying *PDR1^G1079E^*(see methods section). S629P and S663P are the most prevalent and R631G and R665G are the least prevalent mutations found among clinical ECR isolates (14, 16, 28).

**Figure 2.**
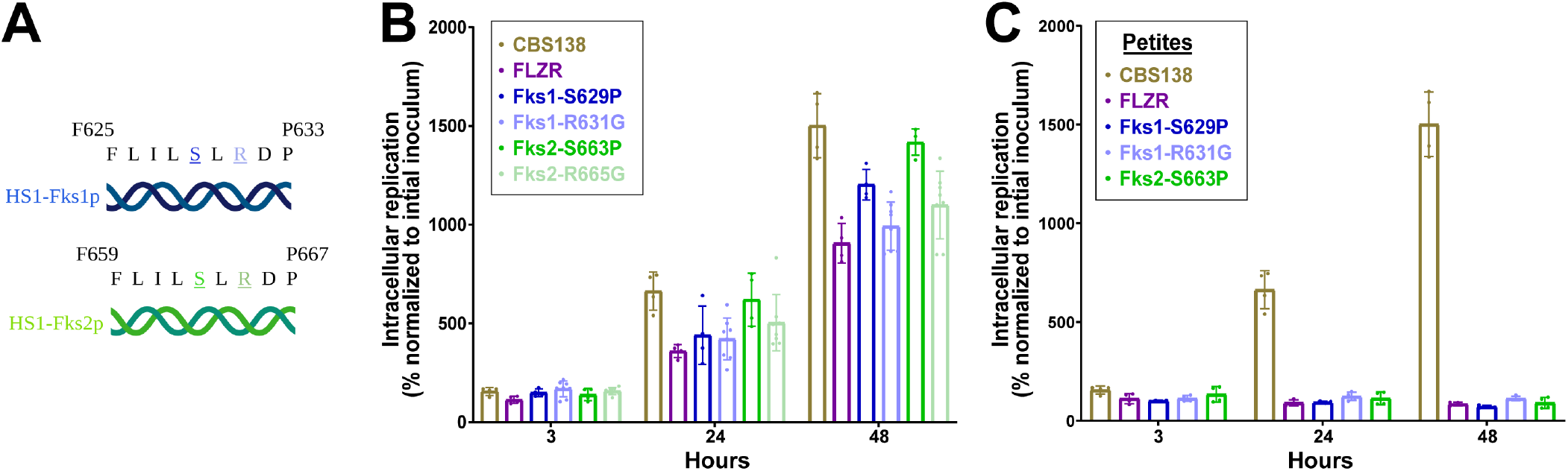
Effects of multidrug resistance on *C. glabrata* survival and replication within macrophages. **A.** Fks1 and Fks2 hot-spot region sequences showing the sites of equivalent amino acid changes in Fks1 and Fks2 representative of clinically prevalent ECR alleles. **B.** Effects of clinically prevalent mutations in HS1 of FKS1 (S629P and R631G) and equivalent mutations in Fks2 (S663P and R665G) introduced into the FLZR parental background. The resulting MDR isolates had a significantly higher IR rate compared to the FLZR parental strain. **C.** Petite MDR isolates carrying the same Fks mutations as in **B** did not show a difference in IR compared to the parental petite isolates. MDR= multidrug resistant, FLZR= Fluconazole resistant, HS1=hotspot 1, IR= Intracellular replication.

The CRISPR-Cas9-generated MDR isolates, the parental FLZR strain, and CBS138 (susceptible wild-type control) were used to infect THP1 macrophages, and IR rates were determined at 3, 6, 24, and 48 hpi. Interestingly, and in line with our expectations, all *fks* mutations could rescue the growth defect of the FLZR parental strain inside the macrophages, albeit to variable degrees (Figure 2C). The extent of this rescue was dependent both on the *FKS* locus and the specific mutation, with mutations in *FKS2*-HS1 showing a higher IR rate compared to their counterparts in *FKS1*-HS1. Interestingly, mutations with a higher clinical prevalence also had a significantly higher IR rates than the less prevalent mutations at the same locus (i.e., S629P and S663P vs. R631G and R665G) (Figure 2B). Of all four MDR mutants, MDR-FKS2^S663P^ had the highest IR rate, comparable to that of the susceptible strain, CBS138.

Petite *C. glabrata* isolates are also known to confer FLZR (25, 26), but we have shown that regardless of origin (clinical or laboratory) such isolates are unable to replicate inside macrophages (https://www.biorxiv.org/content/10.1101/2023.06.15.545195v1). Thus, we asked whether the same clinically relevant *FKS1* and *FKS2* mutations, introduced into petite isolates by CRISPR-Cas9, could rescue their intracellular growth defect. The resulting MDR petite isolates had the same intracellular growth defect as their parental FLZR petite isolates (Figure 2C), consistent with the fact that the petite phenotype mainly stems from defective mitochondria and cannot be restored by introducing echinocandin resistance.

Overall, these observations suggest that the intracellular growth defect of FLZR isolates carrying GOF mutations in *PDR1*, but not mitochondrial dysfunction (petite), can be rescued by *fks* mutations conferring ECR, reinforcing our previous observations that isogenic and clinical MDR strains have higher IR rates than FLZR strains.

### FLZR isolates are outcompeted by MDR and susceptible isolates inside macrophages

In a complementary approach to determine relative fitness of various drug-resistant isolates, we carried out intracellular competition assays. To facilitate these assays, genes encoding green or red fluorescent proteins (GFP or RFP) were stably integrated into the genomes of various isolates to generate constitutively GFP- or RFP-expressing strains. Next, macrophages were infected with ∼50:50 mixtures of two isolates expressing different colors. The macrophages were lysed 3 and 24 hpi, and the proportion of each isolate was determined by flow cytometry. We generated GFP-expressing CBS138 and MDR-*FKS2^S663P^* isolates and RFP-expressing MDR-*FKS1^S629P^* and FLZR isolates. As such, our intracellular competition assayed the competitive intracellular fitness of susceptible *vs.* FLZR, susceptible *vs.* MDR-*FKS1^S629P^*, susceptible *vs.* MDR-*FKS2^S663P^*, MDR-*FKS2^S663P^ vs.* FLZR, and MDR-*FKS2^S663P^ vs.* MDR-FKS1^S629P^.

Our previous studies indicated that GFP and RFP expression do not impose a fitness cost on the IR rate of CBS138 (https://www.biorxiv.org/content/10.1101/2023.06.15.545195v1). Interestingly, the FLZR isolate was outcompeted by the MDR-*FKS2^S663P^*and susceptible counterparts at 24 hpi (Figures 3A and 3B). Moreover, MDR-*FKS2^S663P^*outcompeted the MDR-*FKS1^S629P^* isolate (Figures 3C). Competition between susceptible *vs.* MDR isolates (Figures 3D and 3E) showed that the MDR-*FKS2^S663P^* isolate better competed with the susceptible isolate, consistent with our mono-infection observations. Collectively, these competition assays reinforced the macrophage infection experiments above and underscored that MDR isolates have a fitness advantage over FLZR isolates and are able to replicate better inside macrophages.

**Figure 3.**
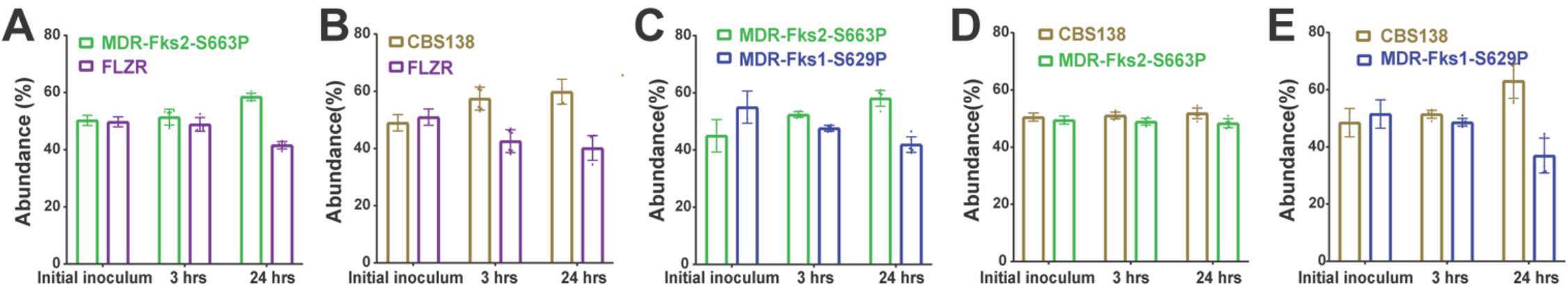
Results of intra-macrophage competition experiments among susceptible and DR *C. glabrata* strains. **A.** MDR-Fks2^S663P^ *C. glabrata* strain outcompeted its FLZR parental strain in macrophages. **B.** Susceptible *C. glabrata* strain outcompeted the FLZR strain in macrophages. **C.** MDR-Fks2^S663P^ *C. glabrata* strain outcompeted the MDR-Fks1^S629P^ strain in macrophages. **D. S**usceptible *C. glabrata* strain had equivalent intra-macrophage fitness with the MDR-Fks2^S663P^ strain. **E.** Susceptible *C. glabrata s*train outcompeted the MDR-Fks1^S629P^ strain in macrophages. MDR= multidrug resistant, FLZR= Fluconazole resistant, HS1=hotspot 1, IR= Intracellular replication.

### Dual-RNAseq analysis

To better understand the host response toward FLZR, MDR, and ECR isolates and *vice versa*, we performed a dual-RNAseq analysis to assess pathogen and host transcriptomes. We selected the pan-susceptible CBS138 and the FLZR isolates, as well as the MDR-*FKS2^S663P^* and MDR-*FKS2^R631G^* isolates since these *fks* mutations had the highest and lowest impact on the intra-macrophage fitness of the parental FLZR isolates, respectively. THP1 macrophages were infected with these isolates and RNA samples were isolated at 3 and 24 hpi and analyzed by RNAseq.

### Transcriptomic responses of *C. glabrata* isolates to macrophage internalization

To get a general overview of the transcriptomics profiles of all studied *C. glabrata* samples we performed a principal component analysis (PCA) (Figure 4A). First, we observed that the analyzed samples clustered predominantly based on interaction with macrophages and on the time point. It is worth mentioning that the factor of time has a stronger impact on *C. glabrata* strains when they grow on RPMI media, as compared to growth within macrophages. Second, the drug susceptibility profiles of the strains had a moderate effect on the overall expression profiles, with susceptible isolates clustering relatively close to FLZR and MDR strains across time points and media.

**Figure 4.**
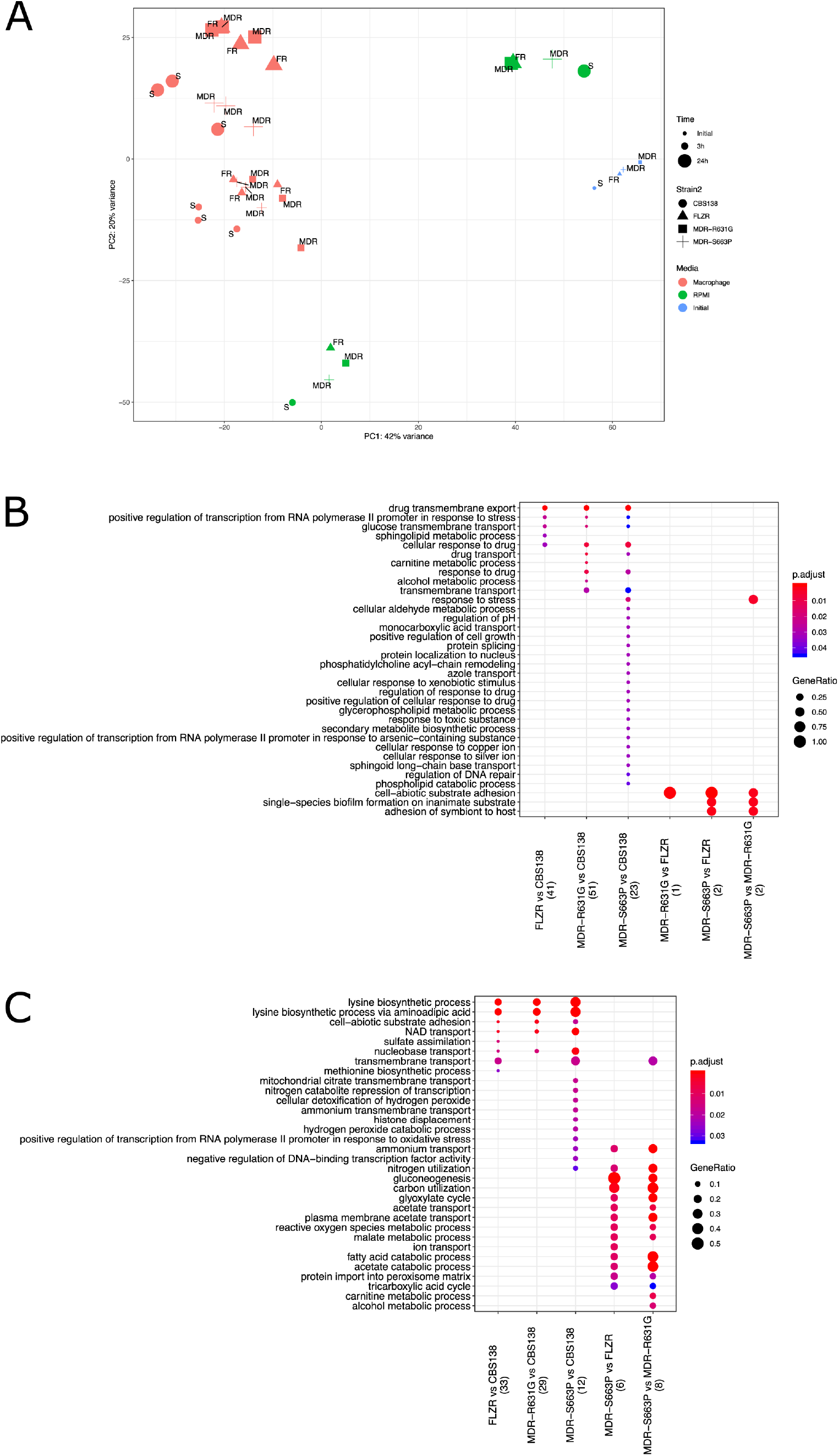
Gene expression changes in susceptible and DR *C. glabrata* strains upon macrophage infection. **A.** Principal Component Analysis (PCA) plot of all studied C. glabrata samples across studied conditions. The plot is based on vst-transformed read count data generated by DESeq2. Labels on the data points correspond to drug susceptibility profiles of each strain: S - Susceptible, FR - Fluconazole Resistant, MDR - Multidrug Resistant. Percentages on PC1 and PC2 axes indicate the total amount of variance described by each axis. **B.** GO term enrichment analysis (category “Biological Process”) of up-regulated genes at a given comparison of C. glabrata strains shown on the X axis. **C.** GO term enrichment analysis (category “Biological Process”) of down-regulated genes at a given comparison of C. glabrata strains shown on the X axis. For (B) and (C) the numbers underneath the comparisons correspond to the “counts” of clusterProfiler (i.e. total number of genes assigned to GO categories). GeneRatio corresponds to the ratio between the number of input genes assigned to a given GO category and “counts”. Only significant (padj<0.05) enrichments are shown. Adjustment of pvalues is done by Benjamini-Hochberg procedure.

We further performed differential gene expression analysis (Supplementary Figure 1) and GO term enrichment of differentially expressed genes (DEGs). Genes with fold-change > 2 or < −2 and adjusted p-value < 0.01 were considered as differentially expressed. The following pairwise comparisons have been performed for the macrophage interaction stage: each drug-resistant strain against the CBS138 (i.e., strain FLZR vs CBS138, MDR-*FKS1^R631G^* vs CBS138, and MDR-*FKS2^S663P^* vs CBS138), multidrug resistant (MDR) strains against fluconazole resistant (FLZR) strain (i.e., MDR-*FKS1^R631G^* vs FLZR and MDR-*FKS2^S663P^* vs FLZR), and comparison between the two MDR strains (i.e., MDR-*FKS2^S663P^* vs MDR-*FKS1^R631G^*). In all cases, we controlled for the effect of time on gene expression levels.

When comparing FLZR and MDR strains, we only observed a few DEGs. For example, only 3 up-regulated and 2 down-regulated genes were identified in the MDR-*FKS1^R631G^* compared to the FLZR strain. All 3 upregulated genes belong to the family of adhesins, resulting in enrichment of the GO term “cell-abiotic substrate adhesion” (Figure 4B) in both comparisons of FLZR and MDR strains. Of note, *C. glabrata* transcriptomic responses revealed that multiple processes important for intracellular and in-host adaptation were significantly downregulated exclusively for MDR-*FKS2^S663P^*, including gluconeogenesis, carbon utilization, glyoxylate cycle, membrane acetate and ion transport, tricarboxylic acid cycle, carnitine metabolic process, etc. (Figure 4C). Therefore, intracellular MDR-*FKS2^S663P^* had a significantly higher transcriptional rewiring when compared to other MDR carrying *FKS1^R631G^*and FLZR isolates.

Interestingly, and in congruence with the PCA plot, FLZR strains and MDR-*FKS1^R631G^* were transcriptionally more similar to each other than the two MDR strains (MDR-*FKS2^S663P^* and MDR-*FKS1^R631G^*), and accordingly, there are numerous enriched GO terms (both up- and down-regulated, Figures 4B and 4C), shared between MDR-*FKS2^S663P^* vs FLZR and MDR-*FKS2^S663P^* vs MDR-*FKS1^R631G^* comparisons. These results are consistent with the observation that the susceptible and MDR*-FKS2^S663P^*as well as FLZR and MDR-*FKS1^6293P^* had similar replication rates inside macrophages.

### Transcriptomic responses of macrophages to infection by the different *C. glabrata* strains

We first performed a PCA analysis to obtain an overall view on the transcriptomes of macrophages infected with the different *C. glabrata* strains (Figure 5A). The results show a clear temporal stratification with distinct clusters of samples at 3 and 24h. Additionally, at 24 hpi the macrophages interacting with CBS138 (drug susceptible) and MDR-*FKS2^S663P^*form a cluster distinct from that of macrophages interacting with other drug-resistant strains, while at 3 hpi all samples show a somewhat uniform transcriptional profile, regardless of the infecting strains.

**Figure 5.**
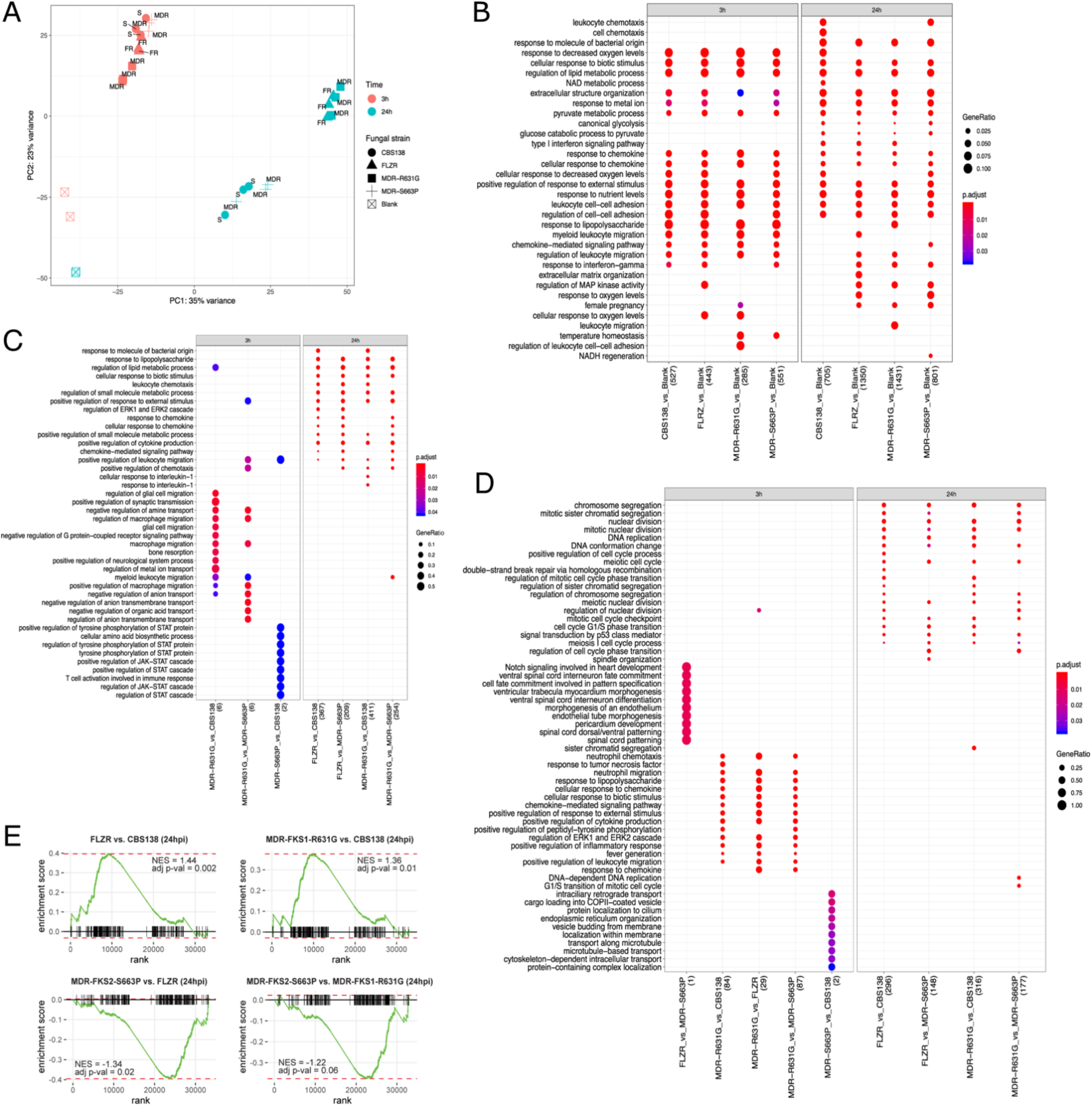
Gene expression changes in macrophages infected with susceptible and DR *C. glabrata* strains. **A.** Principal Component Analysis (PCA) plot of all studied macrophage samples. The plot is based on vst-transformed read count data generated by DESeq2. Labels on the data points correspond to drug susceptibility profiles of infecting *C. glabrata* strains: S - Susceptible, FR - Fluconazole Resistant, MDR - Multidrug Resistant. Percentages on PC1 and PC2 axes indicate the total amount of variance described by each axis. **B.** GO term enrichment analysis (category “Biological Process”) of up-regulated genes of macrophages infected with *C. glabrata* strains (as depicted on X axis) compared to uninfected macrophages. **C.** GO term enrichment categories (when available) of up-regulated genes of macrophages infected with different *C. glabrata* strains (see the X axis for specific comparisons). **D.** GO term enrichment categories (when available) of down-regulated genes of macrophages infected with different *C. glabrata* strains (see the X axis for specific comparisons). For (B), (C) and (D) the numbers underneath the comparisons correspond to the “counts” of clusterProfiler (i.e., total number of genes assigned to GO categories). GeneRatio corresponds to the ratio between the number of input genes assigned to a given GO category and “counts”. Only significant (padj<0.05) enrichments are shown. Adjustment of p-values is done by Benjamini-Hochberg procedure. **E.** Transcripts related to “classically” activated macrophages show significantly enriched expression in the macrophages infected with FLZR and MDR-FKS1^R631G^ strains. Plots depict enrichment of the “classically” activated macrophage transcriptional module (61) for macrophages infected with the indicated strains at 24hpi. Normalized enrichment score (NES) and adjusted P values are shown in the inset.

To further disentangle the transcriptomic differences in macrophage gene expression due to different infecting strains, we performed differential gene expression and GO enrichment analysis comparing uninfected macrophages with macrophages interacting with fungal strains at both time points of infection. This analysis (Figure 5B for up-regulated terms, and Supplementary Figure 2 for down-regulated terms) shows that the transcriptomic profiles of macrophages interacting with each *C. glabrata* strain are largely similar, with the majority of GO term enrichments shared across comparisons.

To get a more detailed representation of the differences in macrophage responses to different infecting strains, especially at 24h, we directly compared macrophages infected with different strains with each other and performed a GO term enrichment analysis (Figures 5C and 5D). Consistent with the PCA plot (Figure 5A), the transcriptional profiles of macrophages infected with CBS138 and MDR-*FKS2^S663P^* were similar to each other and produced a limited number of GO enrichments only at 3h, and the same was true for macrophages infected with FLZR and MDR*-FKS1^R631G^* strains. Furthermore, at the 24h time point the comparisons between macrophage responses to FLZR or MDR-*FKS1^R631G^* vs. CBS138 produced similar GO enrichments to those resulting from the comparisons of FLZR or MDR-*FKS1^R631G^* vs. MDR-*FKS2^S663P^*. This observation underscores that macrophages respond similarly to strains CBS138 and MDR-*FKS2^S663P^* and differently to FLZR and MDR-*FKS1^R631G^*. Although in the light of the susceptibility profiles these results may seem unexpected, it reflects the similar IR rates between MDR-*FKS2^S663P^* and CBS138 strains. On the other hand, and unlike MDR-*FKS2^S663P^*, MDR-*FKS1^R631G^* only marginally increased the defective IR rate of the FLZR strain. In sum, we observed that the more similar the IR rate, the more similar the macrophage transcriptomic responses are.

A similar pattern was observed for both up- and down-regulated genes (Figures 5C and 5D, respectively). Interestingly, macrophages infected with FLZR and MDR-*FKS1^R631G^* at 24 hpi upregulated GO terms related to chemotaxis, lipid metabolism, cytokine production, among others, when compared to macrophages infected with CBS138 and MDR-*FKS2^S663P^*(Figure 5C).

In the same comparison, macrophages infected with FLZR and MDR-*FKS1 ^R631G^* strains predominantly downregulated processes related to cell division, such as chromosome segregation, nuclear division, mitotic cell-cycle checkpoint, among others.

Gene set enrichment analysis (GSEA) of infected macrophages at 24h timepoint revealed that pathways such as “inflammatory response”, “TNFA signaling” were significantly enriched in the macrophages infected with FLZR and MDR-*FKS1^R631G^* when compared to infections with strains CBS138 or MDR-*FKS2^S663P^* (Supplementary Table 3, FGSEA). Furthermore, GSEA using macrophage transcriptional modules of “classically” activated M1 or “alternatively” activated M2 macrophages revealed the M1 transcriptional module to be significantly enriched in the FLZR- and MDR-*FKS1^R631G^*-infected macrophages (Figure 5E).

Collectively, major differences observed at 24 hpi are associated with GO terms related to chemotaxis, lipid metabolism, cytokine production, among others, while down-regulated pathways are mainly related to cell division.

### FLZR and MDR *C. glabrata* isolates are outcompeted by the susceptible parent strain during gastrointestinal tract colonization

As the Gastrointestinal (GI) tract is thought to be a major reservoir for the development of drug-resistant *C. glabrata* mutants (29, 30), we set out to investigate the fitness of FLZR, MDR-*FKS2*^S663P^ and susceptible isolates using a well-established GI tract mouse model (30). Gut colonization was induced via oral gavage, which contained a ∼50:50 mixture of two isolates, one of which constitutively expressed GFP. Fecal samples collected at days 1, 3, 5, and 7 were plated on YPD agar. To differentiate between fluorescent and non-fluorescent colonies, plates were subjected to imaging by a Typhoon Laser Scanner (Cytiva). Because Typhoon is unable to distinguish between RFP and GFP, this competition assay only included GFP-expressing and non-fluorescent isolates.

We previously showed that GFP-expressing CBS138 are slightly less fit in the GI tract than their non-fluorescent counterparts (https://www.biorxiv.org/content/10.1101/2023.06.15.545195v1). Nonetheless, our GI-tract colonization revealed that a GFP-expressing susceptible isolate readily outcompeted non-fluorescent FLZR (Figure 6A) and MDR-*FKS2*^S663P^ isolates (Figure 6B). Surprisingly, the GI-tract competition also showed that a non-fluorescent FLZR isolate readily outcompeted a GFP-expressing MDR-*FKS2*^S663P^, which had previously shown high fitness inside macrophages (Figure 6C). Altogether, these results indicated that susceptible isolates readily outcompeted both FLZR and MDR isolates and that FLZR isolates may be more fit than the MDR isolates in the context of the GI tract.

**Figure 6.**
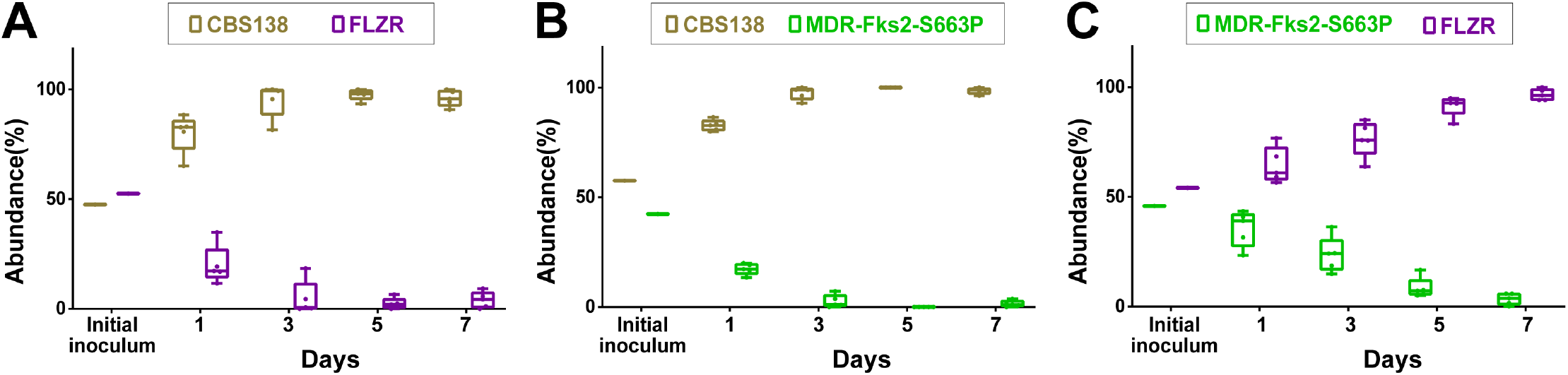
Results of GI tract colonization competition experiments among susceptible and DR *C. glabrata* strains. **A.** Susceptible strain readily outcompeted the FLZR strain. **B.** Susceptible strain outcompeted the MDR-Fks2^S663P^ strain. **C.** FLZR isolate outcompeted the MDR-Fks2^S663P^ strain. MDR=multidrug resistant, FLZR=Fluconazole resistant, GFP=green fluorescent protein.

### The fitness of FLZR and MDR strains varies depending on the infection niche during systemic infection

To investigate the fitness of the FLZR and MDR-FKS2^S663P^ compared to their susceptible counterparts in the context of systemic infection, we used a systemic infection mouse model of *C. glabrata* (30). Systemic infection was induced via the rhino-orbital route using *C. glabrata* cell suspensions containing a mixture of two isolates. Each competition arm included 12 mice (36 mice for three competitions); 4 mice from each arm were sacrificed at each time point. Kidney and spleen collected at days 1, 4, and 7 were homogenized, plated on YPD agar, and the proportions of fluorescent and non-fluorescent colonies were determined using Typhoon.

We previously showed that the fitness cost of the GFP expressing CBS138 varied depending on the organ (https://www.biorxiv.org/content/10.1101/2023.06.15.545195v1), where no fitness cost was observed in the spleen, whereas a fitness cost was observed in the kidney only at day 7. In general, the outcome of the competition in kidney reflected our GI-tract colonization observations, with susceptible isolate outcompeting both FLZR and MDR-FKS2^S663P^ isolates and FLZR isolate outcompeting MDR-FKS2^S663P^ (Figures 6A-6C). However, unlike in the kidney, we found that both FLZR and MDR-FKS2^S663P^ isolates showed similar fitness in spleen (Figure 6D-6F). Collectively, these results indicate that the trajectory of the persistence of drug-resistant and susceptible cells during systemic infection varies depending on the organ. Susceptible and FLZR isolates have an advantage over the MDR isolates in the kidney, whereas spleen is a more permissive anatomical niche, which favors the retention of both resistant and resistant cells.

**Figure 7.**
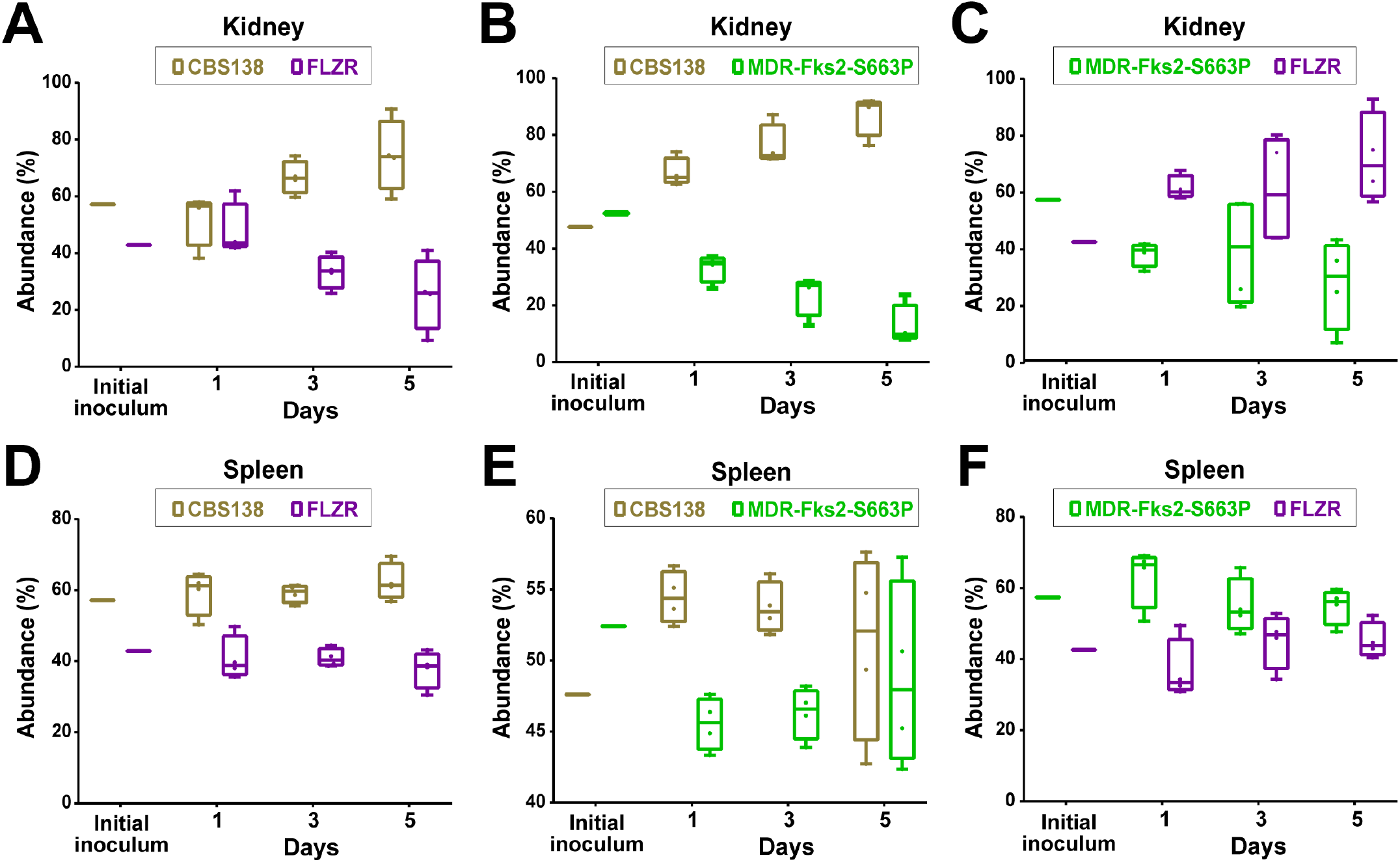
Results of *in vivo* systemic infection competition experiments. **A.** Susceptible isolate readily outcompeted the FLZR isolate in the kidney. **B.** Susceptible isolate readily outcompeted the MDR-Fks2_S663P_ isolate in the kidney. **C.** MDRFks2_S663P_ was outcompeted by the FLZR isolate in the kidney. **D.** FLZR isolate showed comparable fitness in the spleen to the susceptible strain. **E.** MDR-Fks2_S663P_ isolate showed comparable fitness in the spleen to the susceptible strain. **F.** MDRFks2_663P_ and FLZR showed similar fitness in the spleen. MDR= multidrug resistant, FLZR= Fluconazole resistant

## Discussion

*C. glabrata* is one of the most prevalent fungal pathogens causing systemic infections in humans and is characterized by a relatively high number of MDR isolates compared to other *Candida* species (31). Although there are many examples of drug-resistant mutations affecting pathogen fitness in the host with important implications for the spread of AMR (1), the fitness cost of MDR in *C. glabrata* remains poorly understood. Herein, we show that MDR *C. glabrata* isolates displayed improved intra-macrophage growth relative to their parental FLZR strains, and that MDR isolates carrying clinically prevalent mutations in the HS1 of *FKS1* and *FKS2* loci also carried a high intra-macrophage fitness. RNAseq analysis revealed that macrophages infected with MDR or FLZR strains induced pro-inflammatory responses and that cellular processes required for in-host adaptations were downregulated in a MDR isolate carrying Fks1^S663P^. Interestingly, although drug-resistant isolates were outcompeted by susceptible counterparts in the GI-tract and the kidney of mouse models, the environment of the spleen was found to be more permissive, wherein drug-resistant strains showed negligible or undetectable fitness loss relative to susceptible isolates. As such, our study supports the notion that persistence and progression of MDR and FLZR *C. glabrata* isolates is favored and fostered in macrophages and macrophage-rich organs, such as the spleen.

Our *in vitro* experiments indicated that MDR and susceptible *C. glabrata* isolates tolerated stresses characteristic of the macrophage/phagosome environment better than FLZR isolates. Consistent with these *in vitro* data, MDR and susceptible isolates had a significantly higher intra-macrophage growth rate compared to FLZR isolates. Interestingly, MDR isolates carrying *FKS1^S629P^* and *FKS2^S663P^*, which are the most common clinically relevant mutations, had a higher IR rate compared with clinically less prevalent mutations, such as *FKS^R631G^* and *FKS2^R665G^* (16, 28). This potentially explains the high prevalence of *FKS1^S629P^* and *FKS2^S663P^* among clinical isolates. Of note, the lower IR rate of FLZR isolates is not simply due to a lower phagocytosis rate, because the number of intracellular *C. glabrata* cells were similar across all phenotypes at 3 hpi and because phagocytosis rates were similar among these clinical isolates. Although the mechanisms underpinning differential intracellular fitness warrant further investigation, recent studies have shown that the FLZR phenotype in *C. glabrata* (28) and *C. lusitaniae* (24) renders such isolates more susceptible to oxidative stresses and that continuous exposure to oxidative stress-inducing agents *in vitro* and *in vivo* selects against the FLZR phenotype (22, 24). On the other hand, echinocandin resistance is accompanied by cell wall changes (32) that may render cell walls more rigid and less permeable for oxidative stress-inducing agents, which may explain how acquisition of ECR mutations rescues the oxidative stress sensitivity and low intra-macrophage fitness of FLZR strains. Of note, petite *C. glabrata* isolates, which are inherently FLZR and do not replicate inside macrophages (26), did not show any improvements in the IR rate after becoming MDR by acquiring mutations in the HS1 of *FKS1* or *FKS2*. This observation is not unexpected because petiteness is a multifaceted phenotype involving defects in mitochondrial function and central metabolism, which cannot be rescued by *FKS* mutations.

Our dual RNAseq revealed that drug-resistant isolates harbored by macrophages profoundly downregulated processes associated with lysine biosynthesis. Although the role of the lysine biosynthesis pathway in the context of *in vivo* fitness has not been investigated in *C. glabrata*, this pathway is critical for normal biofilm formation in *C. albicans* (33), whereby lysine auxotroph isolates showed altered biofilm structure (34). Indeed, robust biofilm structures are central to colonization of various host niches (35, 36). Accordingly, it is plausible that lysine biosynthesis deficiencies of drug-resistant *C. glabrata* isolates may contribute to their being readily outcompeted by susceptible parental strains during gut colonization and systemic infection involving the kidney. Additionally, macrophages harboring drug-resistant fungi but not their susceptible counterparts had a proinflammatory transcriptomic profile, which may also further explain why drug-resistant isolates were readily outcompeted by susceptible counterparts. Intriguingly, the spleen appeared to be more permissive for retention and persistence of drug-resistant isolates, suggesting that macrophage-rich organs may serve as a viable reservoir for the emergence, retention, or progression of drug-resistant *C. glabrata* cells, especially in the absence of selective pressures imposed by antifungal drugs. Therefore, the applicability and success of restricted antifungal treatment for patients infected with drug-resistant *C. glabrata* depends on the niche inhabited by the infecting cells.

Our RNAseq data also revealed that the MDR-*FKS2^S663P^*isolate additionally and exclusively downregulated various cellular process, including acetate metabolism and transport, glyoxylate cycle, fatty acid catabolism, and protein transport through membrane among the others, which are critical for in-host adaptation (37–39). These observations may explain why the MDR isolate carrying *FKS2^S663P^* was the least fit phenotype in the context of gut colonization and in the kidney during systemic infection. Collectively, our dual-RNAseq and multiple *in vivo* mouse models shed light on the fitness cost associated with drug resistant *C. glabrata* isolates and suggest that such fitness costs could be offset in macrophage-rich niches, which may also act as a reservoir for the emergence and enrichment of drug resistant cells (40).

Intriguingly, RNAseq analysis also revealed that macrophages infected with susceptible and MDR-*FKS2^S663P^* isolates upregulated processes associated with cell cycle progression and mitosis. Although macrophages are considered to be terminally differentiated host cells with limited replicative abilities (41), several recent studies have shown that tissue resident macrophages (TRM) as well as monocyte-derived macrophages (MDM) have the self-renewal capacity (41–46). TRM self-renewal has been implicated during steady state, development, and pathogen challenges, especially with influenza virus (42) and helminth infections (41). Additionally, macrophages infected with the *C. albicans* yeast form, but not with the hyphal form, also have been shown to undergo replication (43). Whether inducing macrophage self-renewal is a protective mechanism employed by the host to contain a highly replicative fungal pathogen or a survival strategy directly/indirectly induced by *C. glabrata* remains uncertain and requires future studies.

It should be noted that clinical drug-resistant isolates often acquire secondary mutations, and perhaps epigenetic modifications, to compensate for the fitness cost associated with antifungal resistance (2, 3). Unlike clinical strains, the isogenic drug-resistant *C. glabrata* isolates tested in this study were generated in the absence of host-relevant constraints. As such, future studies using drug-resistant isolates obtained from various mouse organs and sequential isolates collected from humans will shed additional light on the fitness costs of clinical isolates.

## Methods

### Growth conditions, *C. glabrata* strains and characterization

*C. glabrata* strains were incubated overnight at 37^°^C. Before macrophage infection and mice model infection/colonization, *C. glabrata* strains were grown in YPD broth overnight and incubated in shaking incubator (150 rpm and 37^°^C).

The microbiological information of the clinical and isogenic *C. glabrata* isolates are listed in Supplementary Tables 1 and 2. Clinical isolates were pooled from a global collection of *C. glabrata* isolates and we included various sequence types. The generation of the isogenic ECR, FLZR, and MDR *C. glabrata* isolates from the susceptible CBS138 were detailed in first section of the results. FLZR isolates were denoted when a given colony harbored fluconazole MIC≥ 64µg/ml and harbored a GOF mutation in *PDR1*. ECR isolates were denoted if a given colony harbored mutations in/outside of the HS1 and HS2 of the *FKS1* and *FKS2* genes, whereas MDR isolates should have contained an additional GOF mutation in *PDR1*. All clinical and isogenic isolates underwent sequencing of genes involved in fluconazole and echinocandin resistance using PCR and sequencing conditions described previously (47) as well as the antifungal susceptibility testing (AFST). AFST followed the Clinical Laboratory Standard Institute protocol (48).

### Growth curve

Overnight grown *C. glabrata* cells were washed 2 times with PBS and desired growth media were inoculated with each strain to reach the optical density (OD) of 0.1. *C. glabrata* isolates were incubated at 37^°^C, unless stated otherwise, in a Tecan Microplate Reader (Infinte 2000 pro, DKSH) and the dynamic growth of *C. glabrata* isolates were followed up for 15 hours.

### Macrophage infection

To investigate the phagocytosis survival of our *C. glabrata* isolates, we used a THP1 macrophage derived from human acute monocyte leukemia cell line (THP1; ATCC; Manassas, VA). THP1 macropahges were grown in RPMI 1640 (Gibco, Fisher Scientific, USA) supplemented 10% heat-inactivated HFBS (Gibco, Fisher Scientific, USA) and 1% penicillin-streptomycin (Gibco, Fisher Scientific, USA). To induce macrophage activation and attachment, THP1 cells were treated with 100 nM phorbol 12-myrisate 13-acetate (PMA, Sigma), one million of treated were seeded into 24-well plates (one million each well) and incubated at 37^°^C in 5% CO_2_ for 48 hrs to induce attachment and differentiation into active macrophages (40) Subsequently, active macrophages were infected with overnight grown *C. glabrata* cells with the multiplicity of infection (MOI) of 1/1 (1 yeast/1 macrophage), plates were centrifuged (200g, 1 minute), and plates incubated at 37^°^C in 5% CO_2_ for 3 hrs. Three hours pi, all the wells were extensively washed with PBS and fresh RPMI was added. Of note, the MOI of 5/1 was used for the competition assays.

To calculate the IR rate, macrophages were lysed with one ml of ice-cold water at 3, 6, 24, and 48 hpi and plated on YPD agar plates. IR rate was calculated by dividing the intracellular CFU over the CFU of the initial inoculum and data were presented as percentage. The RPMI collected at 3hrs was plated on YPD plates using which the phagocytosis rate was determined. Phagocytosis rate was defined as the supernatant CFU over the initial inoculum CFU and the values were subtracted from 100.

### Generation of MDR isolates using CRISPR-Cas9

MDR isolates were generated from an FLZR isolate (G2B, Supplementary Table 2). We selected 2 mutations in HS1-Fks1, S629P and R631G, and 2 mutations in HS1-Fks2, S663P and R665G. The codons underlying these mutations were adopted from previous studies (14, 47). To introduce each mutation, we used two overlapping ultramer primers in which the codon the desired mutation was introduced. Of note, the PAM site was also mutated using redundant codon sequences to prevent CRISPR-Cas9 cut of the desired amplicons. For each mutation, we carried out two PCR using forward primer and reverse ultramer primer as well as the forward ultramer primer and the reverse primer (Supplementary Table 4). Subsequently, these two PCR products were fused together using short forward and reverse primers, which were sequenced following PCR product purification. Each fused PCR product should carry two mutations, a non-synonymous mutation conferring ECR and a silent mutation in PAM site.

Competent *C. glabrata* cells were prepared using Frozen-EZ Yeast Transformation Kit (Zymo Research) and transformation followed an electroporation-based protocol described previously (49) and gRNAs listed in Supplementary Table 4. The transformants were incubated in fresh RPMI for 2 hours, followed by spreading them on YPD plates containing 0.125µg/ml of micafungin and incubated for one week in 37^°^C. Positive colonies were subjected to PCR using diagnostic primers (Supplementary Table 4) and subjected to sequencing. ECR colonies should contain the two previously described mutations.

### Macrophage damage assay

To measure the extent of damage incurred by *C. glabrata* isolates to macrophages, we measured the level of lactate dehydrogenase using a commercial kit (Sigma) (50). Briefly, macrophages infected with the MOI of 5/1 were extensively washed with PBS 3 hpi, followed by adding fresh RPMI and incubation in CO_2_ incubator at 37^°^C for another 21hrs. After 24 hours, supernatant samples were collected and LDH was determined as described previously (50). The OD value of each replicate was subtracted from that of the background control (uninfected macrophages) and the corrected value was divided by that of high control (uninfected macrophages treated with 0.25% of Triton X-100 for 3 minutes). The corrected normalized values were presented as percentage.

### Flow cytometry

Macrophages were infected with the MOI of 5/1 and non-adherent yeast cells were removed by extensive PBS washing and adding fresh RPMI 3 hpi. *C. glabrata* cells collected from macrophage at designated time-points were subjected to flow cytometry and 50,000 events were recorded for each replicate. PR rate was determined by subjecting 3hrs supernatant to flow cytometry. The data obtained were analyzed by FlowJo software v10.6.1 (BD Biosciences).

### RNA extraction

Macrophages infected with the MOI of 5/1 were extensively washed 3 hpi and fresh RPMI was added to wells to be further incubated at 37^°^C. After extensive PBS wash at each step, macrophages were subjected to a manual RNA extraction protocol described elsewhere. The RNA samples were treated with RNase free-DNase and further purified using RNeasy kit (QIAGEN) per manufacturer’s instruction. The integrity and quantity of RNA samples were confirmed by running RNA samples in 1.5% agarose gel and NanoDrop (ThermoFisher), respectively.

### RNAseq

RNA isolation was performed as described previously (https://www.biorxiv.org/content/10.1101/2023.06.15.545195v1).

We used FastQC v. 0.11.8 (https://www.bioinformatics.babraham.ac.uk/projects/fastqc/) and MultiQC v. 1.1 (51) to perform quality control of raw sequencing data. Read trimming was performed with Trimmomatic v. 0.36 (52) with the parameters <TruSeq adapters: 2:30:10 LEADING:3 TRAILING:3 SLIDINGWINDOW:4:3 MINLEN:50>.

Rea mapping and quantification was done using splice junction-sensitive read mapper STAR v. 2.7.10a (53) with default parameters. For samples comprising exclusively either fungal or human RNA, reads were mapped to the corresponding reference genomes. For samples containing RNA from both organisms, reads were mapped to the concatenated reference genomes. For human data, we used the novel T2T CHM13v2.0 Telomere-to-Telomere genome assembly (54) genome annotations from the NCBI (last accessed on 12 May 2022). We added the human mitochondrial genome of GRCh38 human genome assembly obtained from the Ensembl database (last accessed on 12 May 2022, (55)). Reference genomes and genome annotations for *C. glabrata* CBS138 were obtained from the Candida Genome Database (CGD, last accessed on 12 May 2022,(56)). Potential read-crossmapping rates (i.e. reads that can map equally well to both human and fungal genomes) were assessed with crossmapper v. 1.1.1(57). Differential gene expression analysis was done using DESeq2 v. 1.26.0 (58). Genes with |log2 fold change (L2FC) | > 1, and adjusted p-value (padj) < 0.01 were considered as differentially expressed. All results of differential gene expression analysis are available upon request. Gene Ontology (GO) term enrichment analysis (category Biological Process) was done by ClusterProfiler v. 3.14.3(59). GO term association tables for *C. glabrata* were obtained from CGD (last accessed on 12 May 2022), whereas for human data we used Genome wide annotation for Human (i.e., org.Hs.eg.db) database v. 3.10.0 to perform GO enrichment tests.

### Gene Set Enrichment Analysis

To define differentially enriched pathways in the infected macrophages, fgsea (https://www.biorxiv.org/content/10.1101/060012v3) was performed using the “Hallmark” Molecular Signature Database pathways (60). The gene sets with adjusted p-values less than 0.05 were depicted in the Supplemental Table. To assess an enrichment of transcripts of the “classically” activated macrophages, the relevant gene list was obtained from Xue et al (61) for performing fgsea.

### Gut colonization mouse model

Our GI-tract colonization mice model included 6-week old female CF1 mice (Chrles River Laboratory) using a previously described protocol (30). The GI-tract commensal bacteria were effectively eradicated by subcutaneous administration of piperacillin-tazobactam (PTZ) starting 2 days prior to infection and continued every day until the end of the experiment (day 7). One hundred µl of cell suspensions containing 1.5×10^8^ mixtures of two isolates were transferred to GI-tract by oral gavage, which was denoted as day 0.

Each competition arm included 5 mice. Fecal samples were collected on 4 time-points, 1-, 3-, 5-, and 7-days post-colonization and resuspended in 500µl of sterile PBS. One hundred microliter of these suspensions were streaked on YPD plates containing PTZ and plates were incubated at 37^°^C for up to 2 days. Subsequently, plates were subjected to Typhoon to visualize the colony color, where dark and light colonies represented GFP-expressing and non-fluorescent *C. glabrata* colonies. Proportion of each isolate was determined by dividing the CFU of a given isolate divided by the total CFU and the proportion was presented as percentage.

### Mouse systemic infections

Our systemic mice infection model used 6-week old CD-1 female mice following a previously established protocol (30). Immunosuppression was induced by administration of 150mg/kg of cyclophosphamide 4 days prior to infection, which was deescalated to 100mg/kg every 3 days afterward. Mice were infected via the rhino-orbital route with 50µl of cell suspensions containing a mixture of two isolates. Each competition arm included 12 mice and 4 mice were euthanized and sacrificed at day 1, day 4, and day 7. One hundred µl of extensively homogenized kidney and spleen samples harvested at designated timepoints were streaked on YPD, which were incubated at 37^°^C for 48hrs. Subsequently, plates were subjected to Typhoon to visualize the colony color, where dark and light colonies represented GFP-expressing and non-fluorescent *C. glabrata* colonies. Proportion of each isolate was determined by dividing the CFU of a given isolate divided by the total CFU and the proportion was presented as percentage.

---

**Supplementary Figure 1.**
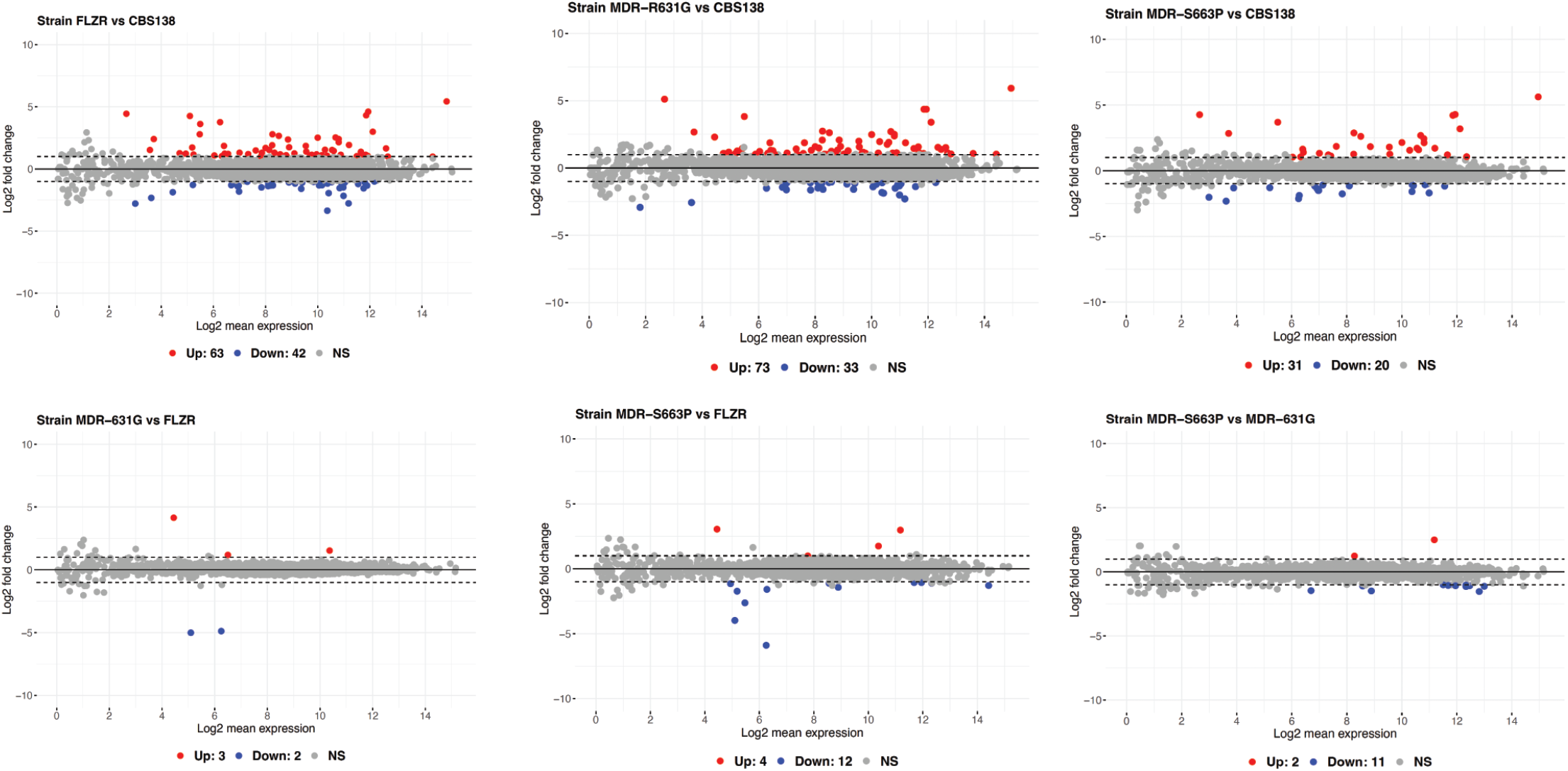
MA-plots displaying the results of differential gene expression analysis. Individual comparisons are specified at the top of each plot. “Up” - up-regulated genes, “Down” - down-regulated genes, “NS” - non-significant. Genes with |log2 fold-change| > 1 and adjusted p-value < 0.01 were considered as differentially expressed.

**Supplementary Figure 2.**
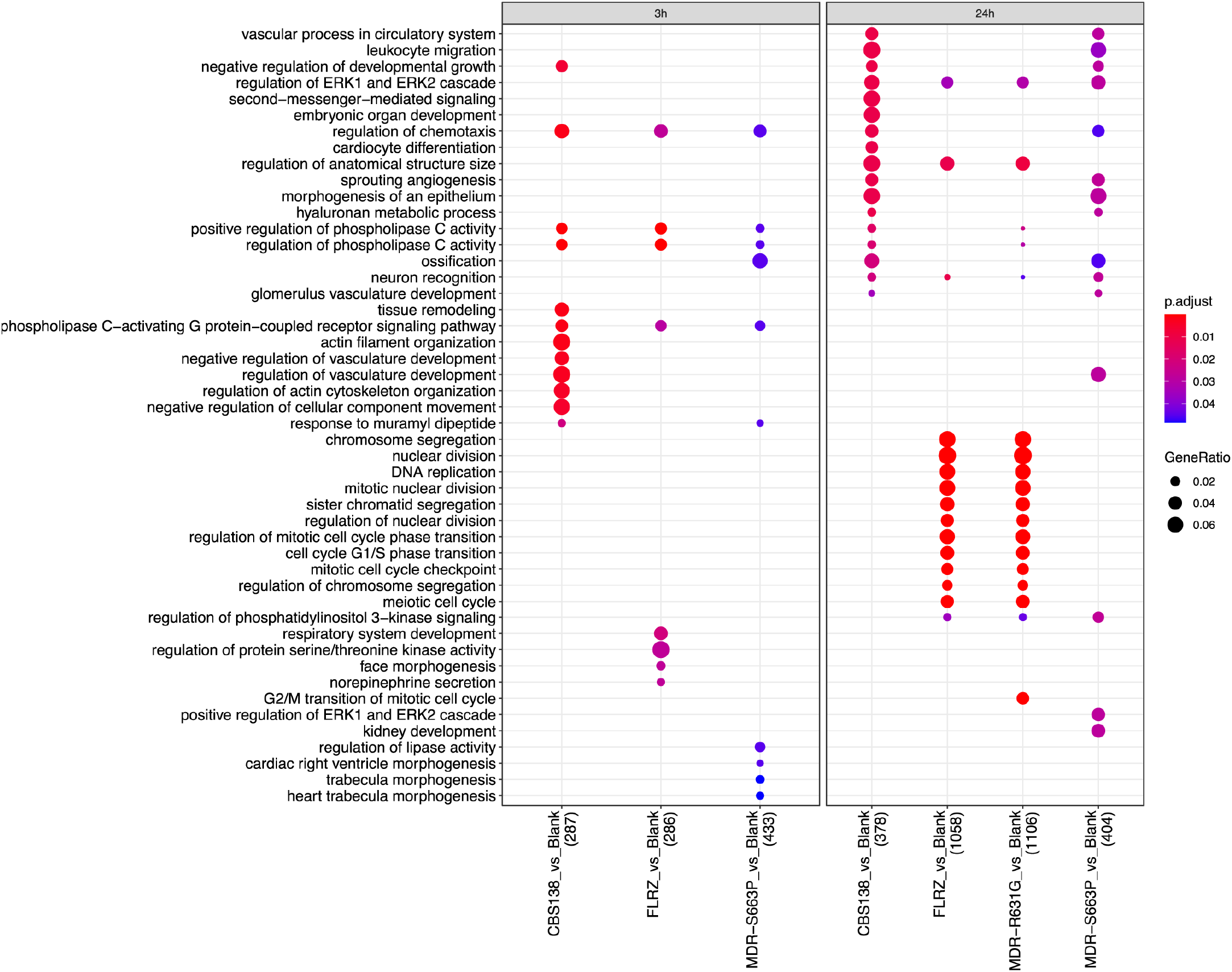
GO term enrichment analysis (category “Biological Process”) of down-regulated genes of macrophages infected with *C. glabrata* strains (as depicted on X axis) compared to uninfected macrophages.

## Notes

### Competing Interest Statement

The authors have declared no competing interest.

